# ATLAS: Analysis Tools for Low-depth and Ancient Samples

**DOI:** 10.1101/105346

**Authors:** Vivian Link, Athanasios Kousathanas, Krishna Veeramah, Christian Sell, Amelie Scheu, Daniel Wegmann

**Affiliations:** Department of Biology, University of Fribourg, Fribourg, 1700, Switzerland; Swiss Institute of Bioinformatics, Fribourg, 1700, Switzerland; Department of Ecology and Evolution, Stony Brook University, 11794, Stony Brook, USA; Institute of Anthropology, Johannes Gutenberg-University, Mainz, 55128, Germany

## Abstract

**Summary:** Post-mortem damage (PMD) obstructs the proper analysis of ancient DNA samples and can currently only be addressed by removing or down-weighting potentially damaged data. Here we present ATLAS, a suite of methods to accurately genotype and estimate genetic diversity from ancient samples, while accounting for PMD. It works directly from raw BAM files and enables the building of complete and customized pipelines for the analysis of ancient and other low-depth samples in a very user-friendly way. Based on simulations we show that, in the presence of PMD, a dedicated pipeline of ATLAS calls genotypes more accurately than the state-of-the-art pipeline of GATK combined with mapDamage 2.0.

**Availability:** ATLAS is an open-source C++ program freely available at https://bitbucket.org/phaentu/atlas.

**Supplementary information:** Supplementary data are available at *Bioinformatics online*.

## 1 Introduction

Ancient genomes provide unique insights into past populations and substantially enhance the inference of demographic and selective events that shaped modern genetic diversity. However, there are two main challenges when genotyping ancient DNA (aDNA): first, only low numbers of unique aDNA fragments remain for sequencing and second, these fragments can be subject to post-mortem DNA damage (PMD). The most common form of PMD is the deamination of cytosin (C), which leads to a C → T transition on the affected and a G → A on the opposite strand after amplification and sequencing (Briggs and Stenzel, 2007). The probability of a C deamination is highest at the ends of the DNA fragments, as these are often single-stranded, and then decays roughly exponentially towards their center (Jónsson *et al*., 2013). As these transitions are artifacts not reflective of the original genome, they must not be considered as variants.

The presence of PMD in the data can be reduced by an enzymatic treatment that cleaves fragments at C’s affected by PMD. However, this technique is restricted to PMD at unmethylated C’s (Briggs *et al*., 2010) and removes precious material. Alternatively, the false calling of variants can be reduced bioinformatically, for instance by trimming reads of their first few nucleotides (e.g. Gamba *et al*., 2014). This leads to a problematically high loss of data, however, if done conservatively. Data loss is much smaller when using mapDamage 2.0 (Jónsson *et al*., 2013), which incorporates the effect of PMD into genotyping pipelines such as Paleomix (Schubert *et al*., 2014) by rescaling base quality scores to additionally reflect the probability of being damaged. But this also leads to a loss of information since PMD is only accounted for indirectly.

We here present the Analysis Tools for Low-depth and Ancient Samples (ATLAS), a collection of statistical methods built upon a dedicated genotyping model that comprehensively accounts for PMD (Hofmanová *et al*., 2016; Kousathanas *et al*., 2017). ATLAS works directly from raw BAM files (Li *et al*., 2009) and contains all necessary methods to accurately genotype and estimate genetic diversity from ancient samples. ATLAS further includes many auxiliary tools to build complete and customized pipelines to work with aDNA or other low-depth samples.

## 2 Methods

### 2.1 Recommended pipeline for single-end data

ATLAS is written in C++, uses the BamTools library (Barnett *et al*., 2011) to parse and write BAM files, and implements many tools, listed in supplementary section 1. Here we present some highlights by outlining the recommended pipeline for single-end sequencing data of ancient samples.

#### Step 1: Classify reads by length

Since PMD rates depend on the distance from fragment ends, ATLAS distinguishes reads spanning the entire fragment, for which these distances are known, from those shorter than their fragment.

**Fig. 1.**
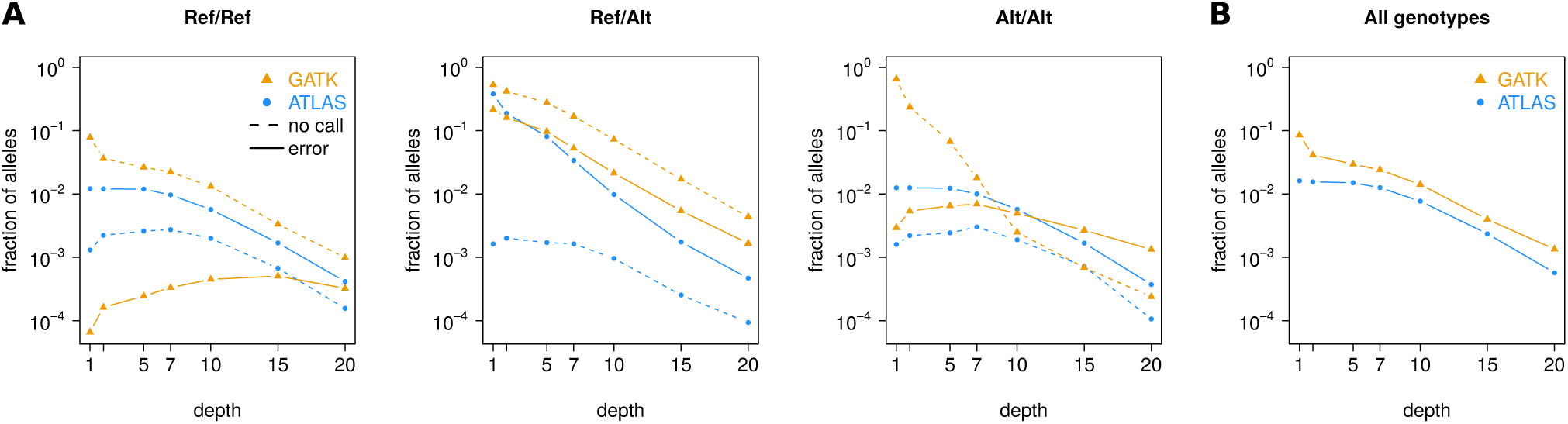
A) Fraction of alleles not (dashed) or wrongly (solid) called at sites with sequencing depth >0 in a simulated chromosome of 10Mbp. The calls are classified according to the underlying true genotype (Ref=reference allele, Alt=alternative allele). B) Fraction of alleles that were not correctly called (either no or a wrong call) with a sequencing depth >0.

#### Step 2: Inferring PMD patterns

ATLAS either infers position-specific PMD patterns, as in MapDamage 2.0 (Jónsson *et al*., 2013), or fits a generalized model of exponential decay (Kousathanas *et al*., 2017), which reduces estimation noise at positions with few PMD observations.

#### Step 3: Recalibrating base quality scores

Base quality scores given by sequencing machines are typically distorted and must be recalibrated. ATLAS offers two recalibration methods we recently developed: a direct extension of Base Quality Score Recalibration (BQSR, DePristo *et al*., 2011) to aDNA (Hofmanová *et al*., 2016) applicable to populations with well characterized polymorphisms, and a reference-free method exploiting haploid or ultra-conserved genomic regions (Kousathanas *et al*., 2017).

#### Step 4: Variant Calling

ATLAS offers three variant callers: a Maximum-Likelihood (MLE) caller similar to GATK (DePristo *et al*., 2011), a Bayesian caller that forms a prior from local diversity and nucleotide composition, and a Bayesian haploid-level caller for very low sequencing depth that identifies the allele with the most evidence to be present using the same prior. For population-level samples, ATLAS further produces individual-specific gVCF files to be analyzed jointly by GATK.

### 2.2 Additional functionalities

For paired-end data ATLAS requires a slightly different pipeline (Supplementary Section 2). Aside from genotype calling, ATLAS also estimates region-specific heterozygosity while accounting for genotype uncertainty (Kousathanas *et al*., 2017), offers tools to reduce modern contamination (similar to PMDS Skoglund *et al*., 2014), and generates input files for PSMC (Li and Durbin, 2011) and BEAGLE (Ayres *et al*., 2012), all of which while accounting for PMD.

## 3 Results

Following Kousathanas *et al*. (2017), we simulated a random diploid chromosome of ten Mbp with heterozygosity *θ* = 0.01 and corresponding ancient single-end sequencing data with errors and PMD at rates commonly observed. We used ATLAS and GATK with comparable parameters (supplementary section 3) to recalibrate base quality scores with BQSR and produce MLE genotype calls. Our GATK pipeline additionally included PMD correction with mapDamage 2.0.

As previously reported (Hwang *et al*., 2015), GATK has an inherent reference bias, resulting in about ten times less genotyping errors at true homozygous reference (Ref/Ref) than homozygous alternative (Alt/Alt) sites, depending on sequencing depth (Fig. 1A). This bias is further accentuated since GATK also less frequently emitted calls at Alt/Alt sites, particularly a low depth, and could not be removed with alternative parameterizations (supplementary section 4). In contrast, ATLAS does not show any such bias (Fig. 1A), and will thus result in higher power to detect true differences between ancient and modern samples.

The two methods also implement different strategies to deal with uncertain genotypes. While ATLAS is designed to emit calls for all sites with data, albeit with low quality scores reflecting that uncertainty, GATK ignores many sites where data is limited. This resulted in slightly higher error rates of ATLAS at lower depth (Fig. 1A), but in a larger fraction of the total genome being called correctly (Fig. 1B). At higher depths, and by accounting for PMD, ATLAS made increasingly less errors than GATK.

## 4 Acknowledgements

We thank Pontus Skoglund for valuable comments on an earlier version of this work, which was supported by the Swiss National Foundation grant 31003A_149920 to DW.

